# Cancer-associated *SMARCAL1* loss-of-function mutations promote alternative lengthening of telomeres and tumorigenesis in telomerase-negative glioblastoma cells

**DOI:** 10.1101/2022.09.19.507319

**Authors:** Heng Liu, Cheng Xu, Bill H. Diplas, Alexandrea Brown, Laura M. Strickland, Jinjie Ling, Roger E. McLendon, Stephen T. Keir, David M. Ashley, Yiping He, Matthew S. Waitkus

## Abstract

Telomere maintenance mechanisms are a hallmark of cancer and are required to enable the replicative immortality of malignant cells. While most cancers activate the enzyme telomerase for telomere maintenance, a subset of cancers (~10-15%) use telomerase-independent mechanisms termed alternative lengthening of telomeres (ALT). ALT is characterized by elevated replication stress at telomeres, telomere synthesis via homology directed-repair mechanisms, and is frequently associated with mutations in the *ATRX* gene. Because ALT is absent in non-malignant proliferating cells, therapeutic strategies targeting ALT-mediated telomere synthesis is an area of significant translational and clinical interest. We previously showed that a subset of adult GBM patients with ATRX-expressing ALT-positive tumors harbored loss-of-function mutations in the *SMARCAL1* gene. SMARCAL1 is an annealing helicase involved in replication fork remodeling and the resolution of replication stress. In this study, we used a patient-derived ALT-positive GBM cell line with native SMARCAL1 deficiency to investigate the role of SMARCAL1 in ALT-mediated telomere synthesis and gliomagenesis in vivo. Our results show that inducible rescue of SMARCAL1 expression suppresses ALT indicators and inhibits de novo telomere synthesis in GBM and osteosarcoma cells, suggesting that SMARCAL1 deficiency plays a functional role in ALT induction in cancers that natively lack SMARCAL1 function. Further, SMARCAL1-deficient ALT-positive cells can be serially propagated in vivo in the absence of detectable telomerase activity, suggesting that the SMARCAL1-deficient ALT phenotype maintains telomeres in a manner that promotes tumorigenesis. In summary, we show that *SMARCAL1* loss-of-function mutations are permissive to ALT and promote gliomagenesis. We also established isogenic model systems that permit the dynamic modulation of ALT activity, which will be valuable for future studies aimed at understanding the molecular mechanisms of ALT and for identifying novel anti-cancer therapeutics that target the ALT phenotype.

## Introduction

Pathological telomere maintenance mechanisms occur in virtually all adult glioblastomas via the activation of telomerase or telomerase-independent mechanisms termed Alternative Lengthening of Telomeres (ALT)^1–3^. Approximately 80% of adult GBM cases harbor single nucleotide mutations in the promoter region of the *TERT* gene (*TERTp*), which leads to the activation of telomerase^1^. A small subset of telomerase-positive *TERT*p-wildtype GBMs (~5%) activate *TERT* expression via chromosomal rearrangements upstream of the *TERT* gene^2^. The remaining cases maintain telomeres through ALT^2^. ALT utilizes homology directed repair (HDR) mechanisms to maintain telomere length and is characterized by frequent *ATRX* loss-of-function mutations, high levels of DNA replication stress, and a remodeling of telomeric chromatin to an epigenetic state that is permissive to homologous recombination^4–9^.

Loss-of-function mutations in the *ATRX* gene are well established to be associated with the ALT phenotype in adult and pediatric GBMs^2,6,10,11^, IDH-mutant astrocytomas^12,13^, high-grade astrocytomas with piloid features^14,15^, brainstem gliomas (including diffuse intrinsic pontine gliomas)^16,17^, and several other cancer types, including sarcomas^11^. Mechanistically, loss of ATRX function is thought to contribute to ALT induction by dysregulating histone H3.3 deposition at telomeres, thus leading to increased DNA replication stress, formation of DNA double-strand breaks (DSBs), and telomere synthesis via break-induced replication^18,19^. Although *ATRX* mutations are closely associated with the ALT mechanism of telomere maintenance, there remains a substantial percentage of ALT-positive tumors that retain intact ATRX function and for which the mechanism of ALT induction and maintenance are not well understood^9^.

We previously reported that a subset of adult *IDH-wildtype/TERTp-wildtype* GBM patients harbor loss-of-function mutations in the *SMARCAL1* (SWI/SNF related, matrix associated, actin dependent regulator of chromatin, subfamily a like 1) gene^2^. *SMARCAL1* encodes an annealing helicase that is recruited to stalled replication forks by replication protein A (RPA)^20^. After being recruited to stalled forks by RPA, a single-stranded DNA binding protein, SMARCAL1 catalyzes fork regression in a manner that promotes fork stabilization and replication re-start, thus preventing deterioration into DNA DSBs^21,22^. In addition to GBM, *SMARCAL1* loss-of-function genetic alterations have been identified in ALT-positive sarcoma cell lines^2,23^ and sarcoma clinical specimens^24^, as well as germline variants associated with pediatric sarcomas and central nervous system tumors^25^.

In the current study, we sought to investigate the functional role of *SMARCAL1* loss-of-function mutations in an ALT-positive cancer cell lines with native SMARCAL1 deficiency. We used a patient-derived GBM cell line, D06MG, to establish the first orthotopic xenograft model of SMARCAL1-deficient ALT and to examine the effects of dynamically rescuing SMARCAL1 activity on the ALT phenotype. We found that SMARCAL1 activity suppresses phenotypic indicators of ALT, such as C-circles, and inhibits the formation of DNA DSBs at ALT-associated PML bodies (APBs) in G2-synchronized cell cultures. Inducible SMARCAL1 expression inhibited de novo telomere synthesis and was sufficient to suppress tumorigenicity in an orthotopic xenograft model. Our results demonstrate that cancer-associated *SMARCAL1* loss-of-function mutations are permissive to ALT-mediated telomere synthesis. Further, these studies established valuable pre-clinical model systems that permit the dynamic modulation of ALT activity, which may be beneficial for ALT-specific drug screening and elucidating the mechanisms of ALT in future studies.

## Results

### A primary SMARCAL1-deficient GBM culture displays stable indicators of ALT activity following in vivo propagation

We first sought to develop a GBM xenograft from a previously established patient-derived cell line that was shown to be ALT-positive and SMARCAL1 deficient^2^. Patient-derived GBM cells were grown in serum-free neural stem cell (NSC) media as neurospheres (D06MG-NSC) and showed stable SMARCAL1 deficiency and maintained expression of ATRX and DAXX over several serial passages (**Fig. 1A**). Immunofluorescence and fluorescence in situ hybridization co-staining (IF-FISH) for PML and TelC respectively showed that D06MG-NSC cells exhibit readily detectable ALT-associated PML bodies (APBs), consistent with a stable ALT phenotype and telomere length maintenance via ALT **(Fig. 1B**).

**Figure. 1.**
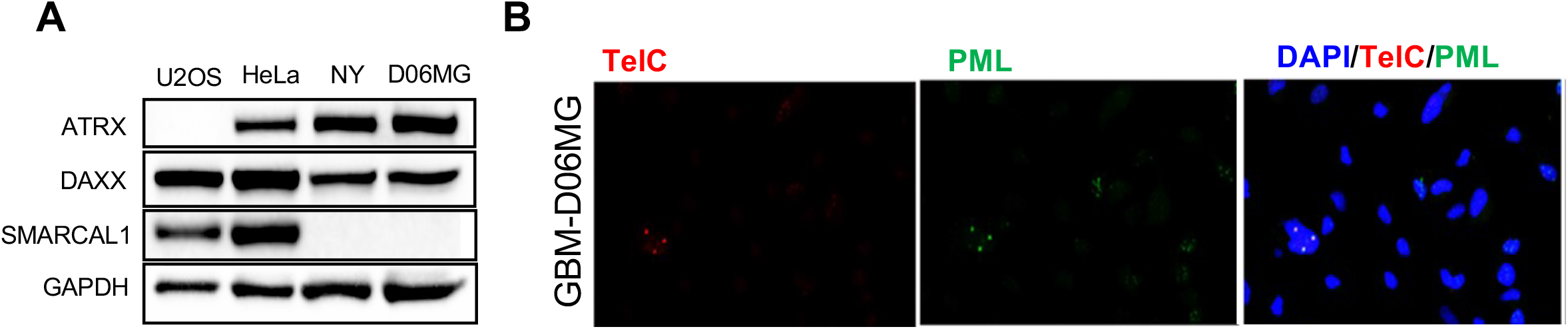
A tumor-derived primary GBM culture is SMARCAL1-deficient and displays ALT indicators. (A) Western blot detection of ATRX, DAXX and SMARCAL1 proteins in total protein extracts. The cancer cell lines U2OS, HeLa, and NY were used for positive and negative controls for the detection of ATRX and SMARCAL1 and compared to expression levels in the primary culture of D06MG cells. (B) Primary GBM culture D06MG was used for detection of telomere and immunofluorescent staining of PML.

We then sought to determine whether D06MG cells were tumorigenic in vivo and to establish xenografts from this cell line. D06MG cells were subcutaneously transplanted into inbred nude mice and allowed to form xenografts over time (**Supplemental Figure 1A**). Subcutaneous tumors formed with a latency of approximately 180 days (**Supplemental Figure 1A, 1B**). Explanted tissue from subcutaneous xenografts was then used to establish a stable cell line under neurosphere growth conditions in cell culture (D06MG-SubQ). In addition, tissue from subcutaneous xenografts was directly passaged forward via intracranial injections into recipient nude mice (**Supplemental Figure 1A**). Orthotopic tumors formed with an approximate latency of 160 days after intracranial transplantation and explanted cells from these tumors were collected and used to establish a cell line in culture (**D06MG-IC**). These studies demonstrated the in vivo tumorigenicity of the D06MG primary culture and established additional xenograft-derived cell lines from the subcutaneous and orthotopic tumors (**Fig. 2A**).

**Figure 2.**
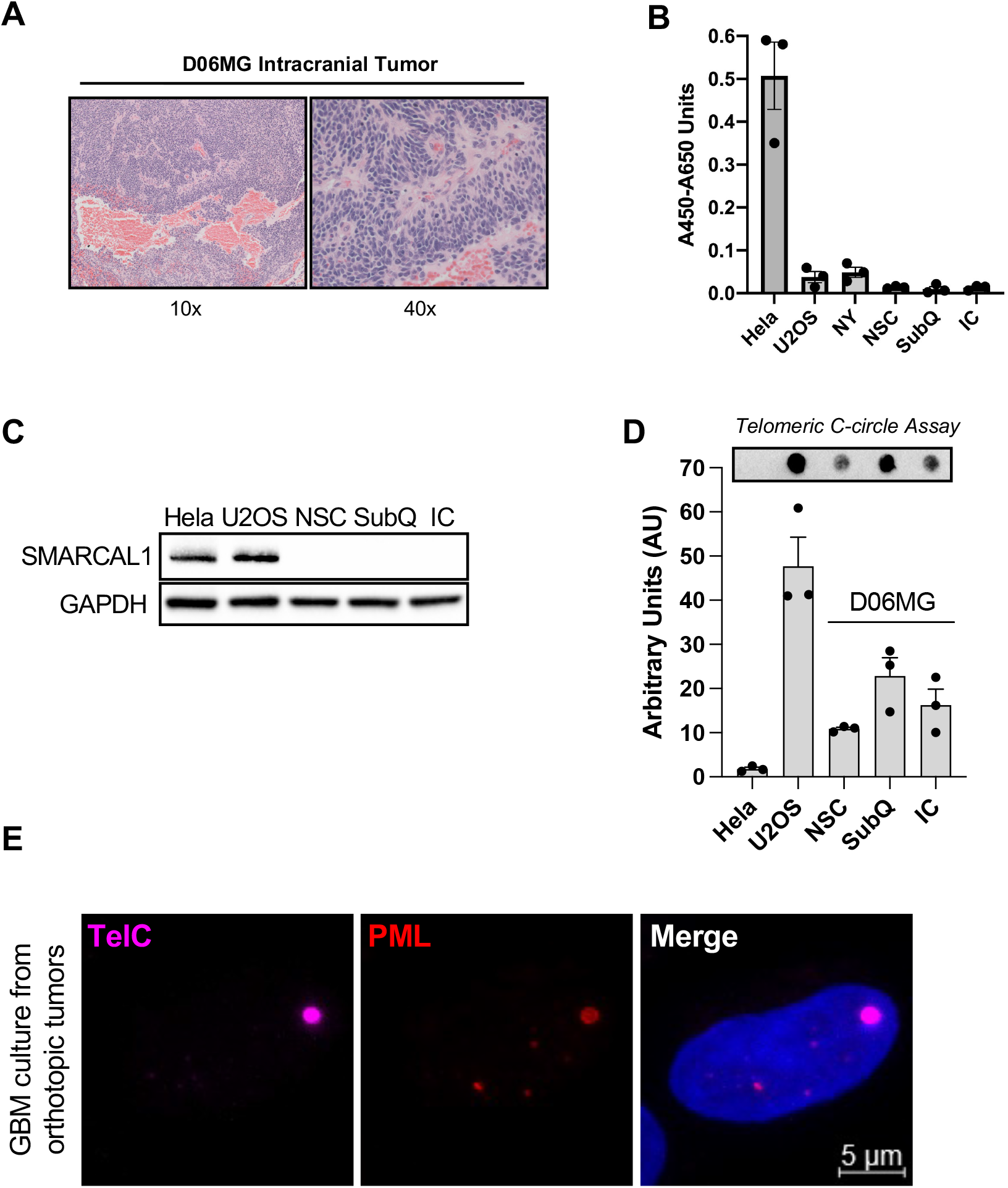
D06MG tumor cells maintain ALT features after in vivo propagation. (A) H&E staining of formalin-fixed paraffin-embedded orthotopic D06MG tumor tissue using 10x and 40x objectives. (B) TeloTAGG assay for the detection of telomerase activity between cell lines. HeLa and U2OS were included as positive and negative controls for telomerase activity, respectively. 3 independent replicates are shown for each condition and the error bars represent the mean +/- standard deviation. (C) Western blot for the detection of SMARCAL1 protein on total protein lysates from D06MG cell lines, as well as positive controls HeLA and U2OS. (D) C-circle assay for the detection of extrachromosomal circular telomeric repeats. 3 independent replicates are shown for each condition and the error bars represent the mean +/- standard deviation. (E) Confocal imaging using a 63x objective of IF-FISH co-staining for telomeric DNA using a TelC-AlexaFlour-647-conjugated TelC probe and an anti-PML antibody detected using an AlexaFluor-594-conjugated secondary antibody and counter-stained with DAPI.

Similar to the parental D06MG-NSC cell line, cell lines obtained from the subcutaneous and orthotopic tumors exhibited low or undetectable telomerase activity (**Fig. 2B**), demonstrating that the intrinsic ALT activity of D06MG is sufficient to maintain telomeres in a manner that promotes tumorigenesis. D06MG-SubQ and D06MG-IC lines were SMARCAL1 deficient (**Fig. 2C**) and displayed ALT indicators, including the presence of extrachromosomal C-circles (**Fig. 2D**) and APBs (**Fig. 2E**), demonstrating that the ALT phenotype is maintained in these cells after in vivo propagation. Because SMARCAL1 is involved in the maintenance of genome integrity and stability^20^, we analyzed the karyotypes of D06MG cells before and after propagation as xenografts in vivo. We found that all D06MG cell lines (prior to and post in vivo growth) displayed mostly diploid karyotypes (**Fig. 3A-B, Supplemental Table 1**). While triploid and tetraploid metaphase spreads were identified in the parental D06MG-NSC culture, D06MG-SubQ and D06MG-IC showed a trend toward a slightly reduced overall number of chromosomes, as well as a lack of triploid or tetraploid cells after in vivo tumorigenesis and propagation (**Fig. 3C**). All metaphase spreads analyzed from both D06MG-SubQ and D06MG-IC showed chromosomal translocations between chromosomes 4 and 22 [(4;22)(q11;q10)] and additional abnormal copies of chromosome 7 (**Fig 3B, Supplemental Table 1**). These distinct alterations were absent in metaphase spreads of D06MG-NSC, suggesting that there was a clonal expansion of a specific tumorigenic subclone following the initial in vivo propagation of D06MG-NSC.

**Figure 3.**
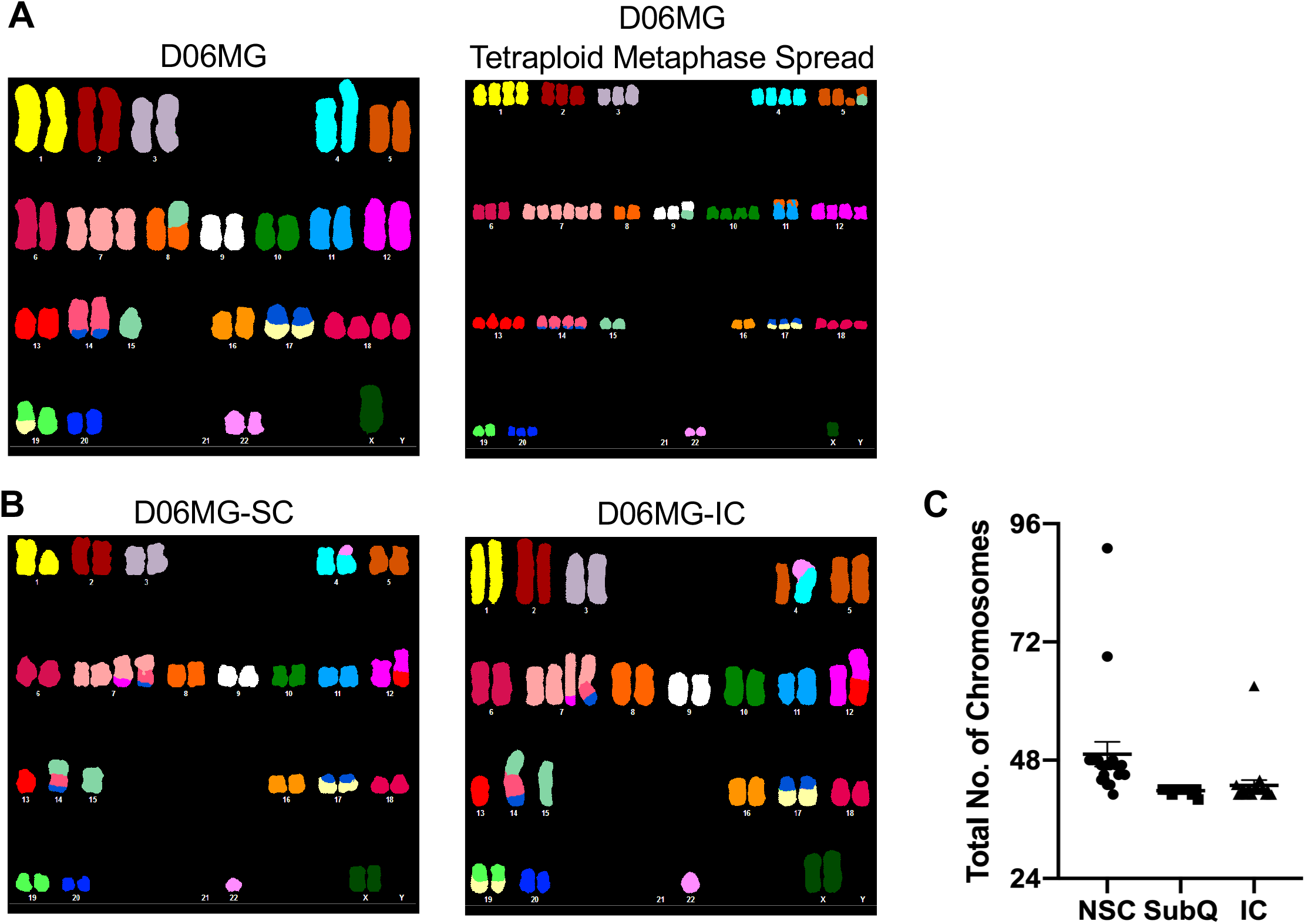
D06MG xenografts are diploid and genomically stable. **(A)** Spectral karyotyping (SKY) analyses of D06MG-MSC primary cell line cultures prior to in vivo propagation. A subset of triploid and tetraploid metaphase spreads were identified, with the majority of cells being diploid. (B) SKY analysis of D06MG cells derived from subcutaneous tumors and orthotopic tumors. (C) Graph showing the total number of chromosomes present in each of the 20 metaphase spreads analyzed per condition.

### Transcriptomic analyses identify gene expression programs associated with tumorigenesis in D06MG xenografts

To determine the gene expression differences associated with the establishment of subcutaneous and intracranial tumors of D06MG, we performed transcriptomic profiling (mRNA-seq) for GBM cells that were maintained in NSC media in vitro (NSC tumor cells), GBM cells derived from the subcutaneous xenografts (D06MG-SubQ), and GBM cells derived from the orthotopic xenografts (D06MG-IC). We noted that in comparison to parental D06MG cells, D06MG-IC and D06MG-SubQ cells displayed remarkably similar pathway-level gene expression changes using the Hallmark pathway analysis^26^. These changes include the modulation of cell cycle gene expression, including the G2M checkpoint, mitotic spindle, the E2F pathway, and the p53 pathway (**Fig. 4A-B, Supplementary Table 2**). Furthermore, D06MG cells undergoing in vivo propagation exhibited higher expression of genes in the hypoxia response pathway, as well genes related to glycolysis, potentially indicating their adaption to in vivo growth (**Fig. 4A-B, Supplementary Table 2**). We next compared the gene expression changes in the D06MG-IC cells relative to the D06MG-SubQ cells. Gene expression changes in the most significantly altered set of genes were those related to the epithelial to mesenchymal transition (EMT) pathway (**Fig. 4C**). While both D06MG-IC and D06MG-SubQ showed decreased expression of EMT genes relative to the parental cell lines, the extent of EMT gene down-regulation was greater in D06MG-IC cells (**Fig. 4C, Supplementary Table 2)**.

**Figure 4.**
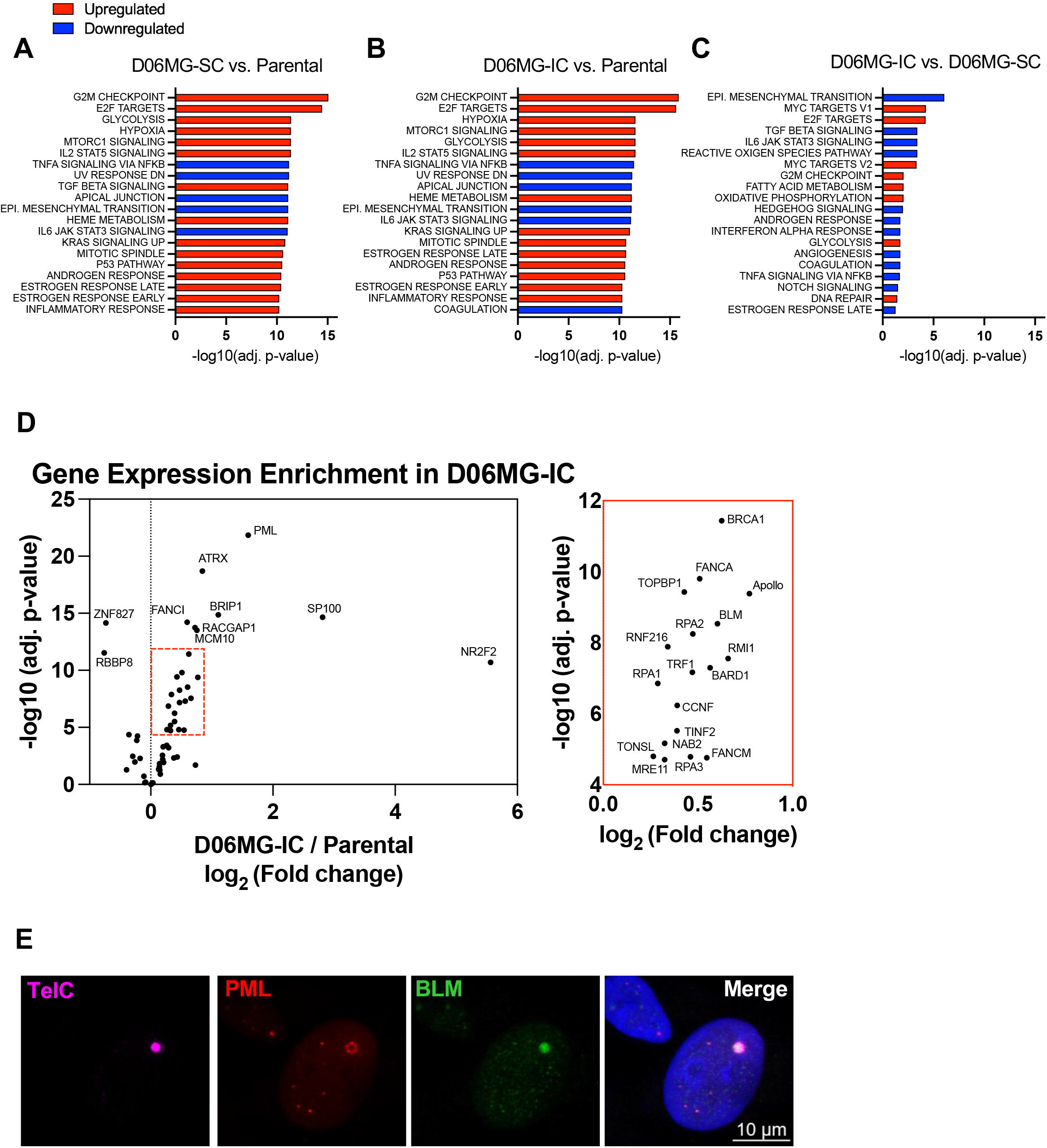
Differential gene expression programs associated with tumorigenesis in D06MG xenografts. **(A)** Analysis of cancer hallmark pathway identified altered pathways in subcutaneous tumor cells versus the initial NSC medium-maintained cells; **(B)** orthotopic tumor cells versus the initial NSC-maintained cells; and **(C)** orthotopic tumor cells versus subcutaneous tumor cells. Pathways were listed by the order of adjust p values. In each figure, a red dash line marks adjusted p value of 0.05. **(D)** Volcano plot of genes encoding proteins localized to ALT telomeres plotted by log2 fold change and adjusted p-value when comparing mRNA expression of D06MG-IC cells relative to the parental cell line. The red dashed box indicates the expanded portion of the plot visualized to the right. Gene identifiers are labelled next to the individual data points. (E) IF-FISH staining of D06MG-IC cells with antibodies to PML and BLM, co-stained with a TelC-647 telomere probe and counter-stained with DAPI.

We then examined differential regulation of genes previously reported to be involved in telomere maintenance or telomere localization in ALT-positive cells. To this end, we used a list of genes encoding for proteins that have been shown to localize to ALT telomeres using proteomic analyses in U2OS cells^27^. We also included *ATRX* because the previously published proteomic analysis was conducted in ATRX-deficient cells, rather than SMARCAL1 deficient cells that have intact ATRX expression (i.e. D06MG). Comparing expression of these ALT telomere-associated genes, we observed that there was a trend toward increased mRNA expression of genes associated with ALT in D06MG-IC cells, including ATRX, PML, FANCM, RPA1/2, and BLM (**Fig. 4D**). While the overwhelming trend was toward increased expression of these ALT-associated genes, there was also a notable decrease in expression of RBBP8 and ZNF827 (**Fig. 4D**). Due to the previously reported roles of BLM at ALT telomeres^28^, we asked whether the observed increase in BLM mRNA expression coincided with BLM localization to APBs in D06MG-IC. Indeed, BLM nuclear foci co-localized with APBs in these cells (**Fig 4E**).

### SMARCAL1 deficiency promotes ALT-mediated telomere synthesis and orthotopic tumorigenesis

We previously reported that SMARCAL1 deficiency was associated with ALT in a subset of adult GBM cases and a chondrosarcoma cell line^2^. In addition, we showed that SMARCAL1 rescue suppressed phenotypic indicators of ALT, including C-circles and ultrabright telomeric foci^2^. We therefore sought to assess the functional role of SMARCAL1 deficiency in promoting ALT. To this end, we utilized a doxycycline-inducible expression system that permits the controlled expression of transgenic SMARCAL1^WT^ or SMARCAL1^R764Q^, a catalytically inactive mutant^2,20,29^ (**Fig. 5A**). Notably, this type of inducible rescue in ATRX-deficient ALT models (e.g. U2OS, SaOS2) has been historically challenging due to the large size of the ATRX protein. On the other hand, the relatively low molecular weight of SMARCAL1 makes the inducible rescue approach feasible and allows for dynamic control of ALT phenotypes (**Fig. 5A**).

**Figure 5.**
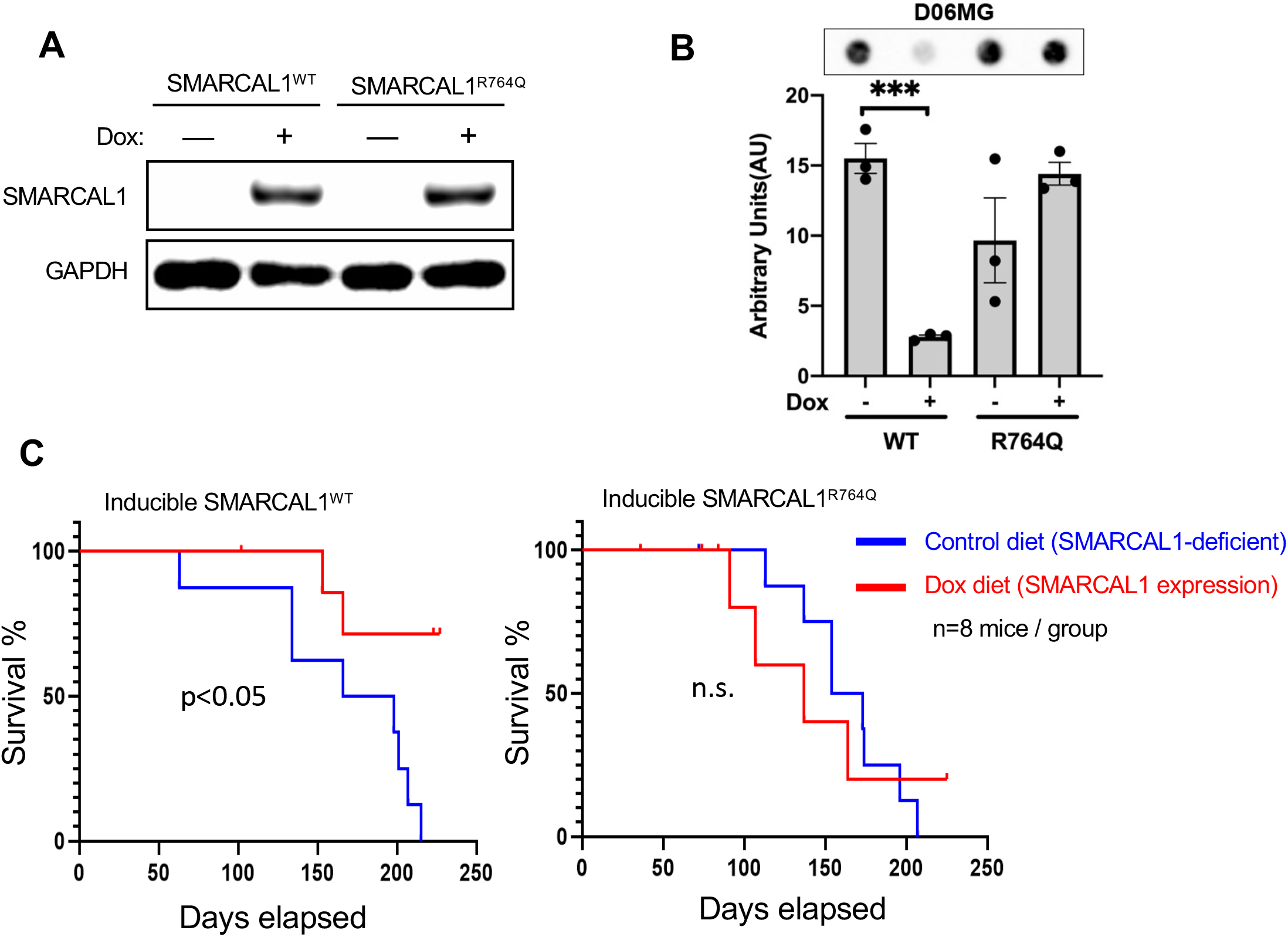
Inducible SMARCAL1 rescue suppresses ALT indicators and in vivo tumorigenesis. (A) Western blot analyses measuring the doxycycline-inducible expression of SMARCAL1 wildtype (SMARCAL1^WT^) protein or an catalytically-inactivate mutant (SMARCAL1^R764Q^) in the primary GBM cultures. (B) C-circle analysis for the primary GBM culture (D06MG) without or with doxycycline-induced expression of SMARCAL1^WT^ or SMARCAL1^R764Q^. SMARCAL1 expression was induced for 5 days and c-circle abundance was measure by dot-blot. Differences between minus-dox or plus-dox conditions was assessed using a student’s t-test. (**C**) D06MG-IC line with inducible SMARCAL1^WT^ (left panel) was orthotopically implanted into mice. 5-days after cell implantation, mice were switched to control or doxycycline-containing chow diet and their survival was determined according to loss of body weight and/or the onset of neurological symptoms. The same experimental framework was used for orthotopic tumors expressing doxycycline inducible SMARCAL1^R764Q^ as a control to account for possible effects of doxycycline chow (right panel). Survival differences between groups was analyzed via the logrank test.

Using this model, we found that inducible SMARCAL1 rescue suppressed the abundance of extrachromosomal telomeric C-circles (**Fig. 5B**), consistent with previous observations and validating the functional rescue of SMARCAL1 activity. We then sought to determine the extent to which SMARCAL1 deficiency regulates tumorigenesis in vivo using orthotopic D06MG-IC tumors with or without doxycycline-induced SMARCAL1 rescue. D06MG-IC cells were transduced with lentiviral particles containing a doxycycline-inducible expression vector for SMARCAL1^WT^ or SMARCAL1^R764Q^. After orthotopic transplantation of these isogenic GBM cell lines into mice, we used a doxycycline-containing diet to induce the expression of SMARCAL1 in vivo. We found that the restoration of SMARCAL1 expression prolonged survival in tumor-bearing mice and decreased the overall penetrance of tumorigenesis (**Fig. 5C**). These SMARCAL1-mediated effects were not observed in the SMARCAL1^R764Q^ mutant condition, suggesting that SMARCAL1 ATPase activity is critical for suppressing tumorigenesis in these cells and ruling out potential effects of doxycycline in vivo (**Fig. 5C**). Collectively, these results demonstrate a functional role of SMARCAL1 deficiency in ALT-mediated telomere synthesis and highlight the importance of this telomere maintenance mechanism in the subset of ALT-positive cancers that exhibit SMARCAL1 loss-of-function.

To gain insight into the mechanism by which SMARCAL1 suppresses orthotopic tumorigenesis in this model, we used inducible SMARCAL1 rescue to investigate the dynamic modulation of ALT indicators and de novo telomere synthesis in D06MG-IC cells. Because ALT utilizes break-induced-replication for maintaining telomere length, we examined the regulation of APBs and the co-localization of γH2AX (a marker of DNA DSBs) in APBs in G2/M-synchronized cells. We found SMARCAL1 expression led to a reduction in APBs that co-localized with γH2AX (**Fig. 6A**). This result suggests that SMARCAL1 expression suppresses replication stress and inhibits the formation of DSBs at telomeres in G2/M phase and is consistent with a role of SMARCAL1 in facilitating replication fork repair and restart to prevent degeneration into DSBs^20,21,30^.

**Figure 6.**
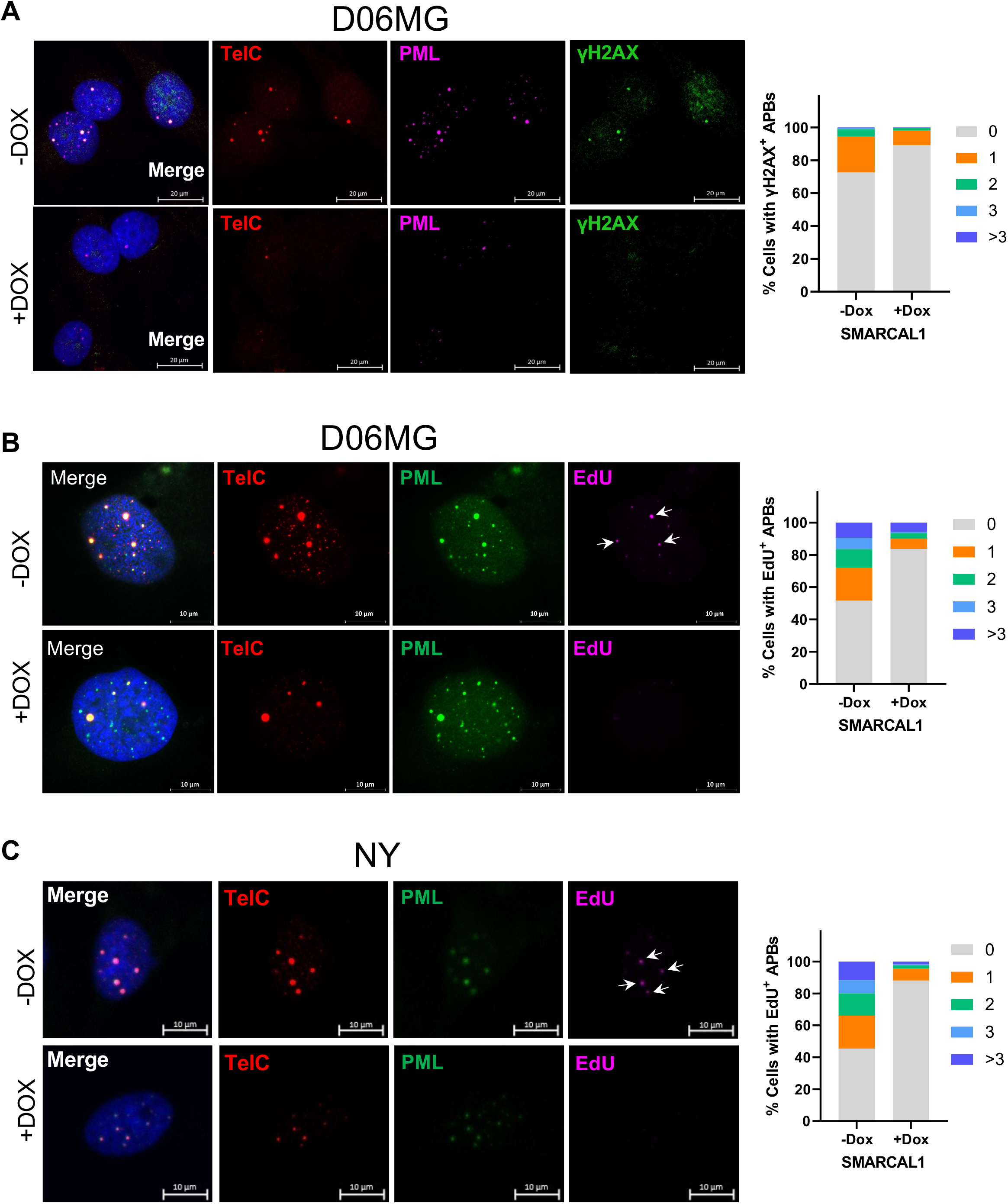
SMARCAL1 activity suppresses telomeric DNA DSBs and de novo telomere synthesis in D06MG cells. (**A**) The primary D06MG-IC culture without or with Dox-inducible SMARCAL1 restoration were used for APB and γH2AX detection and representative images were shown. Cells were seeded in a chamberslide and allowed to adhere before SMARCAL1 expression was induced with doxycycline (1 ug/ml). 5 days after SMARCAL1 induction, cells were fixed and processed for IF-FISH with a TelC probe, anti-γH2AX antibody, and anti-PML antibody. Quantification of APBs that displayed positive γH2AX IF staining. Data represent the proportion of individual nuclei that exhibit APBs with co-staining of γH2AX. **(B)** D06MG-IC cells without or with doxycycline-induced SMARCAL1 expression were pulse-labelled with EdU (2 hours) before processing for telomeric foci detection (FISH) and immunofluorescent detection of nuclear PML bodies with an anti-PML antibody. Co-localization of TelC foci and PML was used to identify APBs, and EdU staining at APBs was used to nascent telomeric DNA synthesis at APBs. (**C**) Experiment performed as in (B) with NY osteosarcoma cells.

We then investigated whether SMARCAL1 loss-of-function is necessary for de novo telomere synthesis in ALT+ cell lines with native SMARCAL1 deficiency. These experiments used D06MG-IC cells and NY cells, an osteosarcoma cell line previously shown to be SMARCAL1-deficient and ALT-positive^23^. Similar to D06MG, NY cells are C-circle-positive at baseline, and SMARCAL1 rescue significantly reduces C-circle abundance in these cells (**Supplemental Figure 2A-B**). We then used EdU pulse-labelling of de novo DNA synthesis in late G2-synchronized D06MG-IC and NY cells and examined the co-localization of newly synthesized DNA with APBs. Doxycycline-inducible rescue of SMARCAL1 expression significantly inhibited the number of EdU+ APBs relative to cells without SMARCAL1 rescue in both D06MG-IC cells and NY cells, demonstrating that SMARCAL1 activity suppresses de novo telomere synthesis in these lines (**Fig. 6B-C**). To validate this finding, we performed a BrdU pulse-labeling experiment to label newly synthesized DNA and subsequently immunoprecipitated BrdU-labelled DNA for telomere probe detection via dot blot. SMARCAL1 rescue significantly inhibited the amount of BrdU-incorporated telomeric c-strand DNA relative to SMARCAL1 deficient controls (**Supplemental Figure 2C**).

## Discussion

In this study, we used a patient-derived ALT-positive GBM cell line with native SMARCAL1 deficiency to establish subcutaneous and orthotopic xenografts, as well as cell lines derived from the xenograft tissue. Using an inducible rescue of SMARCAL1 expression, we found that SMARCAL1 activity suppresses ALT indicators and inhibits de novo telomere synthesis (**Fig. 6A-C, Supplemental Figure 2C**), suggesting that SMARCAL1 deficiency plays a functional role in ALT induction in cancers that natively lack SMARCAL1 function.

Our results establish SMARCAL1 as an additional gene associated with onset of the ALT phenotype. In cancer, loss-of-function genetic alterations in *ATRX* or *DAXX*, which encode histone H3.3 chaperone proteins, are well established to play a role in ALT-mediated telomere maintenance in cancers from several tissue types that are enriched for ALT-positive tumors^7,11^. In addition, cell-based studies have shown that depletion of another histone chaperone, ASF1, leads to rapid induction of ALT and suppression of telomerase activity^31^. It is notable that ATRX, DAXX, and ASF1 all function as histone chaperones, while SMARCAL1 functions as an annealing helicase that is recruited to stalled replication forks with lagging strand gaps by the single-stranded binding protein RPA^20,32,33^. In this way, it remains unclear whether the mechanisms by which SMARCAL1 loss-of-function leads to ALT are similar to, or distinct from, the mechanisms of ALT induction caused by perturbations in histone deposition. However, based on published studies and our results, we speculate that the common feature of proteins whose loss-of-function leads to ALT is an ability to promote replication fork stability within difficult-to-replicate DNA sequences (i.e. telomeres), thereby preventing stalled replication forks from deteriorating into DNA DSBs^19,34–36^. We propose that in the absence of SMARCAL1 activity, there is an increased frequency of stalled replication forks within telomeric DNA that collapse into single and double-stranded DNA breaks and that this increase in telomeric DSBs functions as a nexus for break-induced replication of telomeres (**Fig. 7**)^8,18,34,37–39^.

**Figure 7.**
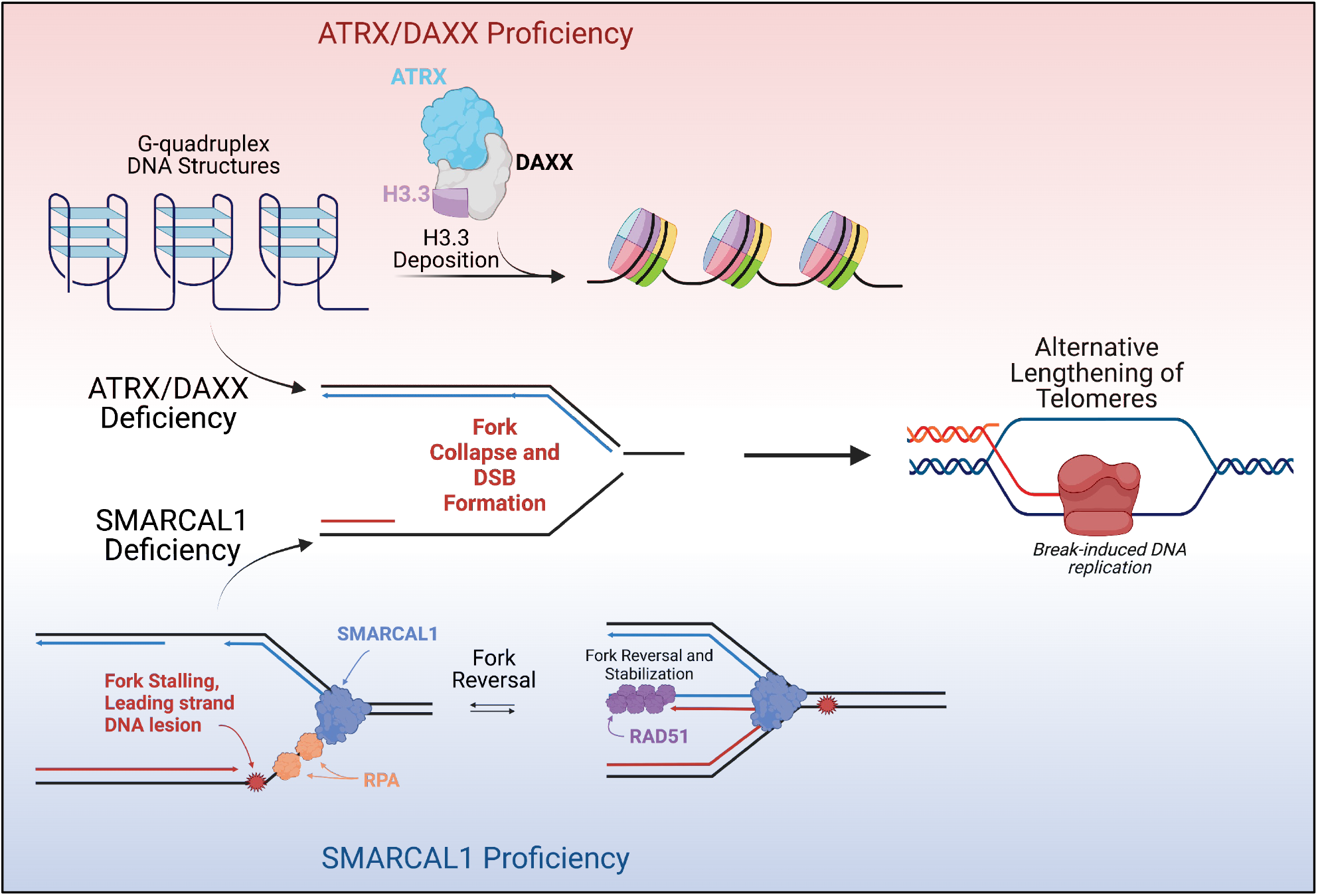
Alternative pathways of ALT induction in human cancers. Cancer-associated loss of function genetic alterations occurring in ATRX/DAXX (top) or SMARCAL1 (bottom) can independently lead to the ALT induction in cancer cells. ATRX/DAXX deficiency compromises the canonical H3.3 chaperone function of these proteins, leading to increased G-quadruplex formation and replication stress at telomeres. SMARCAL1 deficiency leads to a loss of replication fork reversal and stabilization, thus leading to degeneration into DSBs, which in turn facilitate break-induced replication of telomeres. Illustration was created with Biorender.com.

Although we have shown that SMARCAL1 deficiency contributes to ALT induction in a subset of *ATRX* wildtype GBMs, SMARCAL1 activity has also been shown to play a role in resolving DNA replication stress and promoting ALT activity in the ATRX-deficient setting. Cox et al. have shown that SMARCAL1 localizes to APBs in ATRX-deficient ALT-positive sarcoma cells^30^. In that study, the investigators showed that SMARCAL1 activity prevents excessive replication fork stalling at ALT telomeres, thereby inhibiting DSB formation and preventing chromosomal fusions^30^. SMARCAL1 may therefore exhibit a dual function in telomere maintenance in the context of ALT. In the ATRX-deficient context (ATRX-deficient ALT), SMARCAL1 activity appears to be critical for maintaining relative genome stability and promoting telomere maintenance^30^. On the other hand, *SMARCAL1* loss-of-function genetic alterations contribute to ALT-mediated telomere maintenance (SMARCAL1-deficient ALT) in tumors and cell lines that express wildtype ATRX^2,23^. Importantly, the molecular mechanisms of ATRX-deficient ALT and SMARCAL1-deficient ALT must be at least partially distinct, as SMARCAL1 activity plays a critical role in resolving replication stress in the ATRX-deficient setting that, by definition, SMARCAL1 cannot perform in the SMARCAL1-deficient ALT context. The extent to which SMARCAL1-deficient ALT and ATRX-deficient ALT exhibit common and unique mechanisms of telomere synthesis and resolution of replication stress will be investigated in future studies.

The availability of patient-derived ATRX-deficient ALT cell lines and xenografts has been historically limited for a number of reasons. First, the majority of ALT-positive adult glioma cases occur in *IDH-mutant* astrocytomas (grades 2-4), which have been challenging to establish as cultured cell lines^40,41^. Those lines that have been established are almost invariably derived from recurrent cases that have progressed to high-grade tumors and are often pre-treated with TMZ and/or radiotherapy^41,42^. Moreover, because *ATRX* encodes a large protein, it is difficult to develop experimental models with rescued ATRX expression in order to dynamically regulate the ALT phenotype. Our D06MG models are therefore unique in that they are ALT-positive GBM cells that are derived from an untreated primary tumor^2^. Further, because SMARCAL1 is a relatively small protein compared to ATRX, inducible rescue of SMARCAL1 expression is feasible in these cells, allowing the dynamic modulation of the ALT phenotype in vitro and in vivo (**Figs. 5 and 6**). We propose that modulating ALT in this isogenic context may be valuable for future studies aimed at further understanding the molecular mechanisms of ALT and for identifying novel anti-cancer therapeutics that target the ALT phenotype.

In summary, our results demonstrate that SMARCAL1-deficiency promotes ALT-mediated telomere maintenance in a manner that supports tumorigenesis in a patient-derived xenograft model. Although the majority of ALT-positive tumors and cell lines are ATRX-deficient, the underlying genetics and mutational status of *SMARCAL1* and *ATRX* should be considered in future pre-clinical and clinical studies that focus on ALT mechanisms and associated drug sensitivities. Finally, SMARCAL1-deficient ALT-positive models may be useful tools for transiently suppressing the ALT phenotype or phenotype-switching (ALT to telomerase) to create isogenic model systems comparing therapeutic vulnerabilities according to telomere maintenance mechanisms^9^.

## Materials and Methods

### Neurosphere Cell culture

Patient-derived neural stem cell cultures were passaged according to previously published methods^42^ using the NeuroCult^™^ NS-A proliferation Kit (Stemcell^™^ Technologies, Catlog#05751) supplemented with 20 ng/ml human recombinant EGF (Stemcell^™^ Technologies, Cat#78006.2) and 20 ng/ml human recombinant bFGF (Stemcell^™^ Technologies, Cat#78134.1). The parental cell line and xenograft-derived lines were authenticated by Duke DNA Analysis facility. The D06MG cell line was established from archived tissue in the Preston Robert Tisch Brain Tumor Center Biorepository as previously described^2^. D06MG P0 archived cells from the Preston Robert Tisch Brain Tumor Center BioRepository (accredited by the College of American Pathologists) with the approval from the Institutional Review Board

### Xenograft generation and passaging

D06MG cells cultured under neural stem cell conditions were dissociated via Accutase digestion and mechanical trituration, filtered through a 70-micron filter, and re-suspended in a methylcellulose PBS solution for injection into recipient mice. For initial subcutaneous xenograft generation, 5×10^6^ cells were injected per animal. For direct passaging of subcutaneous tumors for intracranial injections, tumor tissue was excised from host mice under sterile conditions in a laminar flow containment hood and homogenized in a modified tissue press. The homogenate was loaded into a repeating Hamilton syringe dispenser and orthotopically injected into the right caudate nucleus of recipient mice under isoflurane anesthesia using a stereotactic apparatus. For intracranial injections from the D06MG-IC cell line, single cell suspension were prepared in methylcellulose/PBS solution as described above and cells were injected into the right caudate nucleus, as previously described^43^.

### Spectral Karyotyping analysis

Spectral Karyotyping preparation and analysis were performed on D06MG cell lines according to established procedures by Herbert Irving Comprehensive Cancer Center (HICCC) Cytogenetics Core, Columbia University. Briefly, cells were cultured attached in laminin coated plates with serum free medium and treated with 0.05 μM colcemid solution for two hours. Medium containing floating cells was collected, spun down and treated with 0.56% KCl hypotonic solution for 20 min at 37 °C. Cells were fixed with methanol/acetic acid (3:1 by volume) fixative solution and dropped to ethanol cleaned slides. After pretreatment with pepsin solution, the slides were denatured and hybridized with chromosome-specific painting probes at 37 °C for 48 hours. Probes were visualized using Cy5-conjugated avidin and Cy5.5-conjugated anti-mouse IgG. Chromosomes were counterstained with 4,6-diamidino-2-phenylindole. 20 metaphases from each cell line were randomly selected and analyzed.

### Immunoblotting

Cells were lysed in RIPA buffer (Millipore Sigma, Cat# R0278) with 1X protease inhibitor cocktail (Cell Signaling Technology, Cat# 5871S). Protein lysates containing ~20 μg total protein were loaded and resolved using (4-12%) NuPAGE Bis-Tris gradient gel. Gels were soaked in protein transfer buffer (48 mM Tris, 39 mM glycine, 20% methanol, 0.0375% SDS) and transferred to a PVDF membrane using a BioRad Mini Tran-Blot transfer cell. After transfer to PVDF membranes, membranes were blocked with Pierce TBST Protein-Free blocking buffer (Cat # 37572) and blotted with antibodies diluted in blocking buffer. Antibodies used included anti-ATRX (Cell Signaling, Cat#14820), anti-DAXX (Cell Signaling, Cat#4533), anti-SMARCAL1 (Cell Signaling, Cat#44717) and anti-GAPDH (Santa Cruz, Cat#sc-47724), γH2AX (Cell Signaling, Cat#9718), PML (Santa Cruz, Cat#sc-966).

### Telomerase PCR ELISA Assay

The activity of telomerase in the cell lines was detected using the TeloTAGGG telomerase PCR ELISA Kit (Roche, Cat# 12013789001). The assay was performed according to the manufacturer’s instructions and repeated in triplicate. 2×10^5^ cultured cells were collected, counted and lysed in 200 μL lysis reagent. 1 μL of cell lysis, negative controls and positive controls were mixed with 25 μL TRAP Reaction buffer. The mixtures were performed a combined primer elongation/amplification reaction following the manufacturer’s standard protocol. The reaction products were hybridized to the microplate precoated with digoxigenin (DIG)-labeled telomere probes, incubated with Anti-DIG-POD antibody and detected by TMB substrate solution. Data was collected by measuring absorbance of the samples at 450 nM with a reference wavelength of 650nM using Tecan i200 Pro plate reader.

### RNA-Seq

Cell line RNA was extracted using Maxwell RSC simplyRNA Cells Kit (Promega Cat# AS1390). Extracted RNA was sent to Novogene (San Diego, CA, USA) for quality control and RNA sequencing. The raw FASTQ data was aligned to reference genome (GRCh38/hg38) using HISAT2^44^. Count reads were mapped to genes using featureCount and DESeq2 was used for counts normalization^45,46^. Limma-voom was used for differential gene analysis^44^. EGSEA was used for gene set enrichment analysis^47^. A false discovery rate (FDR) cut-off of 0.05 of differentially expressed genes was used for the calculation of Significance Ccore and Regulation Direction.

### Immunofluorescence-FISH co-staining

Cells were grown on 3-well chamber slides (Ibidi) to sub-confluence. The cells were fixed for 10 minutes with 4% formaldehyde diluted in PBS and were then incubated with primary antibodies in blocking buffer (1 mg/mL BSA, 3% goat serum, 0.1% Triton X-100, 1 mM EDTA) overnight at 4 °C. After primary antibody incubation, the slides were incubated with goat secondary antibodies against rabbit or mouse IgG, conjugated with AlexaFluor 488 or 594 (ThermoFisher, 1:500). The cells were then rinsed with PBS and fixed with 2% formaldehyde for 10 minutes at room temperature. Following a series of sequential dehydration steps (70%, 95%, and 100% ethanol), cells were incubated with PNA probes (each 1:1000) TelC-Cy3 (PNA Bio, Cat# F1002) or TelC-AlexaFlour-647 (PNA Bio, #F1013) in hybridizing solution, denatured at 70 °C for 10 minutes on a ThermoBrite instrument, and then incubated at room temperature for 2 hours or overnight at 4 °C in a humidity chamber. Slides were then washed with 70% formamide, 10 mM Tris-HCl, PBS, stained with 4’,6-diamidino-2-phenylindole (DAPI), and then cover-slipped and sealed. Slides were imaged on a Zeiss 880 upright/inverted confocal microscope and a Zeiss 780 upright confocal microscope, and images were analyzed using Zen software. For quantitation of foci and co-localization of IF-stained foci with telomeres, the Focinator program was used^48^. Antibodies used included γH2AX (Cell Signaling, Catalog # 9718), PML (Santa Cruz, Catalog # sc-966), and BLM (Santa Cruz, Cat#365753).

### Nascent telomere DNA synthesis labeling and pull down

To track telomere DNA synthesis in the telomeres of ALT positive cells, the ALT telomere DNA synthesis assay (ATSA assay) was performed as previously described^36^. Inducible rescue cell lines were cultured with a treatment of doxycycline (0.5ug/ml) for five days. Dox treated cells and control cells were seeded into chamber slide and synchronized to late G2 phase using Thymidine and CDK1 inhibitor, RO-3306 two-step synchronization. The cells were pulse-labeled with 10 μM 5-ethynyl-2’-deoxyuridine (EdU) or 10 μM 5-bromo-2’-deoxyuridine (BrdU) for 2 hours. Cells were then permeabilized and fixed with 4% formaldehyde in PBS. EdU incorporated cells were stained by Click-iT chemistry (Thermo Fisher) to detect the incorporated EdU, followed by IF-FISH staining using a TelC-Cy3 probe and antibodies against PML, EdU and γ-H2AX. Stained cells were then imaged using a Zeiss 880 confocal microscope. As a separate method for quantifying the amount of newly synthesized telomeric DNA after arresting in G2, cells were pulsed with BrdU for 2 hours, after which cells were lysed for genomic DNA isolation using a Promega Maxwell, RSC Cultured Cells DNA kit. After sonication, nascent telomere DNA was immunoprecipitated using an anti-BrdU antibody (BD Biosciences, Cat#347580). Immunoprecipitated DNA was dot-blotted onto a nylon membrane and detected by biotin-labeled telomere probe and quantified using Bio-Rad densitometry software.

### Mouse Intracranial injection

Glioma stem-like cells were transplanted into the right caudate nucleus of athymic nude mice (1×10^5^ cells per injection, N=8 per group). Mice were placed in a Stoelting stereotactic injection device under isoflurane-induced anesthesia. An incision was made in the midline of the scalp over the position of the frontal cortex, and bregma was located visually. Cells were injected 2mm right and 3.5 mm deep relative to Bregma using a 25-gauge needle, and the needle was raised ~.5 mm. After one minute, the syringe was slowly removed to prevent efflux. The site of injection was then sealed with bone wax, Marcaine was administered as a local anesthetic, and the incision was sealed with surgical glue. Animals were then monitored for the onset of neurological symptoms as evidenced by indicators such as inability to ambulate, agonal breathing, head tilting or doming, or a 20% loss of body weight, and then euthanized. Differences in survival of tumor-bearing mice between conditions were evaluated using the log-rank test.

### Statistical Analyses

For analyzing differences between doxycycline treated or untreated cells, a student’s t-test was used. For c-circle assays with >2 groups, a one-way analysis of variance was used with correction for multiple comparisons by the Bonferroni method. Survival analyses for in vivo animal studies was performed using the log-rank test. For gene expression analyses via RNA-seq, a cut-off of FDR 0.05 of differentially expressed genes was used for the calculation of Significance Score and Regulation Direction. For figures and figure legends in the manuscript, the following annotation are used to denote varying thresholds of statistical significance: *P < 0.05. **P < 0.01. ****P < 0.0001. ns = non-significant.

## Acknowledgements

We thank Drs. Lisa Cameron and Yasheng Gao of the Duke Light Microscopy Core Facility for imaging support and expertise, Drs. So Young Kim, Sufeng Li, and Vidya Seshadri of the Duke Functional genomics Core for their help with generating constructs used in the study, Dr. Wenxia Jiang of the Herbert Irving Comprehensive Cancer Center (HICCC) Cytogenetics Core, Columbia University, for cytogenetic analysis, and Dr. Hai Yan for his guidance and resources during the early stages of this work. We thank the staff of the Cancer Center Isolation Facility (CCIF), Duke Cancer Institute, for assistance with the in vivo experiments. This work was supported by The Preston Robert Tisch Brain Tumor Center and NIH grant 1K22CA258965-01A1 awarded to M.S.W.

## Supplemental Figure Legends

**Supplemental Figure 1.**
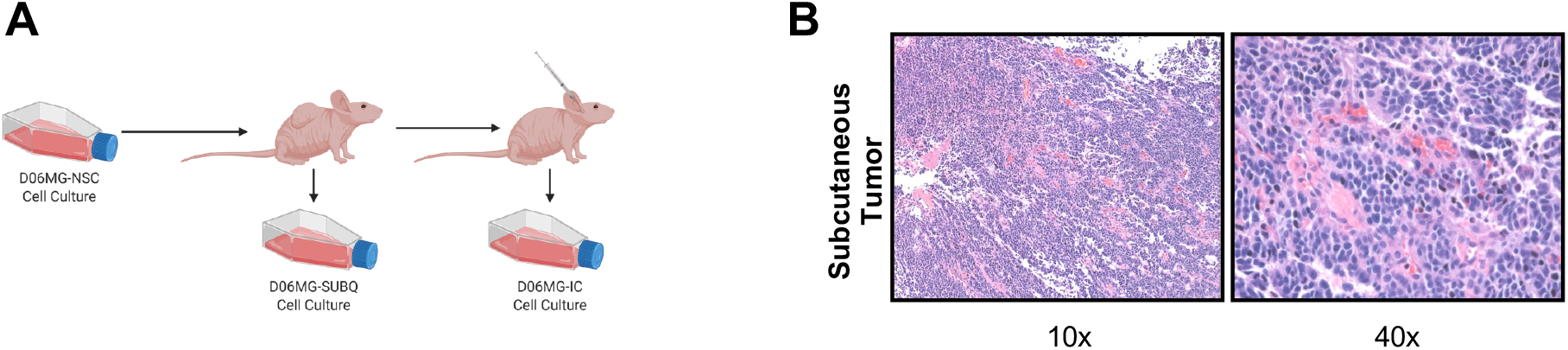
Alternative pathways of ALT induction in human cancers. (A) Schematic representation of the experimental design for serial implantations and propagation of xenografts. Subcutaneous and orthotopic tumors derived from the primary GBM culture were established and cell lines maintained from xenograft tissue. D06MG cells were implanted subcutaneously into nude mice and allowed to establish xenografts. Cells from subcutaneous xenografts were passaged forward in mice to generate orthotopic tumors, as well as explanted into cell culture to establish a stable cell line after in vivo propagation. (B) H&E staining of subcutaneous D06MG tumors using 10x and 40x objectives.

**Supplemental Figure 2.**
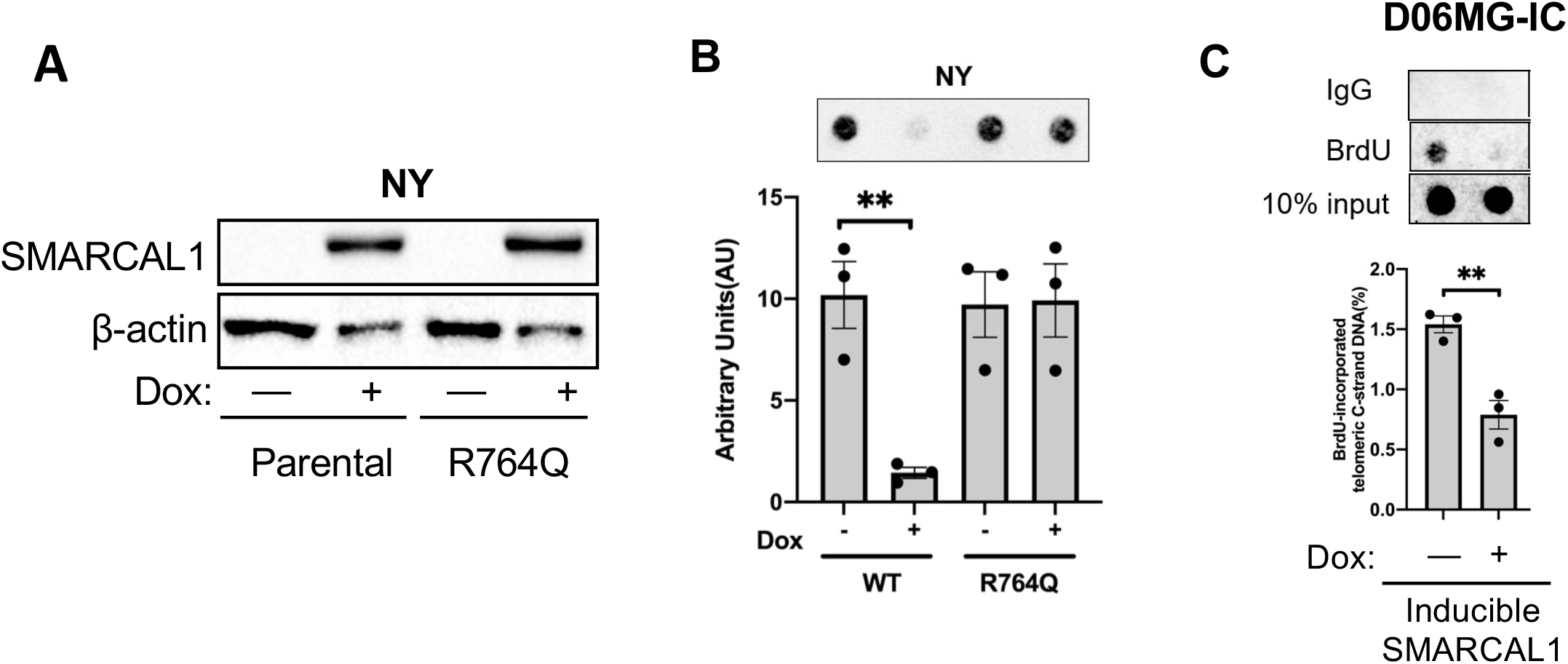
Validation studies of SMARCAL1-mediated regulation of de novo telomere synthesis. (A) Western blot analysis of parental NY cells or NY cells transduced with a doxycycline inducible SMARCAL1^R764Q^ expression vector. Inducible rescue of SMARCAL1 expression was validated 5 days after induction with 1 ug/ml doxycycline. (B) C-circle assay performed using NY cells with or without inducible expression of SMARCAL1^WT^ or SMARCAL1^R764Q^. SMARCAL1 expression was induced for 5 days with 1 ug/ml doxycycline and c-circle abundance was assayed via rolling circle amplification and dot-blot. Differences between minus-dox or plus-dox conditions was assessed using a student’s t-test. (C) The primary GBM cells without or with the Doxycycline-induced SMARCAL1 expression (.5 ug/ml) were labelled with BrdU (2-hours) before being used for telomere pull down followed by detection of telomeric DNA via dot-blot using a TelC probe. Differences between minus-dox or plus-dox conditions was assessed using a student’s t-test.

## Supplemental Tables

**Table S1. Spectral Karyotyping Analyses of D06MG patient-derived GBM cells**

**Table S2. Transcriptomic analyses of D06MG patient-derived GBM cells**

## Author Contributions

- **Conceptualization**: Heng Liu, Bill Diplas, Matthew Waitkus
- **Data curation**: Heng Liu, Yiping He, Stephen Keir, Matthew Waitkus
- **Funding acquisition:** David Ashley, Matthew Waitkus
- **Investigation:** Heng Liu, Cheng Xu, Alexandrea Brown, Laura Strickland, Jinjie Ling, Stephen Keir, Matthew Waitkus
- **Methodology**: Heng Liu, Cheng Xu, Bill Diplas, Stephen Keir, David Ashley, Matthew Waitkus
- **Resources**: Stephen Keir, Roger McLendon, David Ashley, Matthew Waitkus
- **Supervision**: Yiping He, Matthew Waitkus
- **Writing – original draft**: Heng Liu, Yiping He, Matthew Waitkus
- **Writing – review & editing**: Heng Liu, Cheng Xu, Bill Diplas, Alexandrea Brown, Laura Strickland, Jinjie Ling, Roger McLendon, Stephen Keir, David Ashley, Yiping He, Matthew Waitkus

## Notes

### Competing Interest Statement

The authors have declared no competing interest.

### Summary of Updates

Updated Figure Legends for Figs 2 and S1.

